# Caveats to deep learning approaches to RNA secondary structure prediction

**DOI:** 10.1101/2021.12.14.472648

**Authors:** Christoph Flamm, Julia Wielach, Michael T. Wolfinger, Stefan Badelt, Ronny Lorenz, Ivo L. Hofacker

## Abstract

Machine learning (ML) and in particular deep learning techniques have gained popularity for predicting structures from biopolymer sequences. An interesting case is the prediction of RNA secondary structures, where well established biophysics based methods exist. The accuracy of these classical methods is limited due to lack of experimental parameters and certain simplifying assumptions and has seen little improvement over the last decade. This makes RNA folding an attractive target for machine learning and consequently several deep learning models have been proposed in recent years. However, for ML approaches to be competitive for de-novo structure prediction, the models must not just demonstrate good phenomenological fits, but be able to learn a (complex) biophysical model. In this contribution we discuss limitations of current approaches, in particular due to biases in the training data. Furthermore, we propose to study capabilities and limitations of ML models by first applying them on synthetic data (obtained from a simplified biophysical model) that can be generated in arbitrary amounts and where all biases can be controlled. We assume that a deep learning model that performs well on these synthetic, would also perform well on real data, and vice versa. We apply this idea by testing several ML models of varying complexity. Finally, we show that the best models are capable of capturing many, but not all, properties of RNA secondary structures. Most severely, the number of predicted base pairs scales quadratically with sequence length, even though a secondary structure can only accommodate a linear number of pairs.

## 1 Introduction

Many RNAs rely on a well defined structure to exert their biological function. Moreover, many RNA functions can be understood without knowledge of the full tertiary structure, relying only on secondary structure, i.e. the pattern of Watson-Crick type base pairs formed when the RNA strand folds back onto itself. Prediction of RNA secondary structure from sequence is therefore a topic of longstanding interest for RNA biology and several computational approaches have been developed for this task. The most common approach is “energy directed” folding, where (in the simplest case) the structure of lowest free energy is predicted. The corresponding energy model is typically the Turner nearest-neighbor model [19], which compiles free energies of small structure motifs (loops) derived from UV melting experiments.

Under some simplifying assumptions, such as neglecting pseudoknots and base triples, the optimal structure can be computed using efficient dynamic programming algorithms that solve the folding problem in 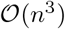 time for a sequence of length *n*. While these algorithms yield an optimal solution given the model, the accuracy achieved on known secondary structures varies widely and averages about 67% in a benchmark accompanying the latest Turner parameter set [11]. While a variety of factors contribute to the inaccuracy of prediction, accuracy has hardly changed in comparison to the previous iteration of energy parameters [12], suggesting that it is the simplifying assumptions of the model, rather than measurement errors in the UV melting experiments, that limits prediction accuracy. It is therefore tempting to forego the simplifying assumptions necessary for dynamic programming and approach the problem using machine learning techniques. Inspired by the recent success of deep learning methods in protein structure prediction, several groups have proposed deep learning methods for the RNA secondary structure prediction problem [3, 18, 17, 8].

A major problem for all deep learning approaches is the limited availability of training data. Even before the recent machine learning boom, several works have attempted to replace or improve the Turner energy parameters by training on a set of known RNA secondary structures [6, 1, 21]. While these works demonstrated that learning energy parameters is feasible, they often reported overly optimistic accuracies. Even if test and training sets do not contain very similar sequences (e.g. with less than 80% identity), this is not sufficient to avoid overtraining. As shown in [15], setting up test/training sets that avoid biases by using *structurally* distinct RNAs leads to a significant drop in accuracy and largely eliminates any advantages of the trained over measured parameters. Thus, ideally test and training sets should be constructed from distinct RNA families.

The currently most used training set machine learning on RNA structures is the bpRNA set [5] which contains over 100000 distinct sequences. While the number of sequences in this set is sufficient to train sophisticated models, the structural diversity of the data set is limited: 55% of the sequences are ribosomal RNAs (rRNAs) from the Comparative RNA website [2]. The next largest data source is the Rfam database [13], providing 43% of sequences. At first glance, this subset seems more diverse, since Rfam release 12.0, used for bpRNA, comprises 2450 RNA families. Again, however, rRNA and tRNAs make up over 90% of the sequences in Rfam 12.0. The dataset is therefore dominated by just four RNA families (three types of rRNA as well as tRNAs) and it seems highly unlikely that it can capture the full variety of the RNA structure space. This is also reflected in the extremely uneven length distribution of sequences in bpRNA, see Fig. S1. We will explore the effect of using a training set with such limited structural diversity in section 5.

When both test and training set are derived from bpRNA, they will exhibit the same biases leading to unrealistically good benchmark results. The MXfold2 paper [17] addressed this problem by generating an additional data set, bpRNAnew, containing only sequences from Rfam families added after the 12.0 release. The bpRNAnew set was also used in the Ufold paper [8] to distinguish between within-family and cross-family performance. In practice, within-family performance is largely irrelevant: structure prediction for sequences belonging to a known family should always proceed by identifying the RNA family and mapping the novel sequence to the consensus structure, e.g. using covariance models and the Infernal software [14]; this is in fact how most of the structures in the bpRNA set were generated. Only sequences that cannot be assigned to a known family should be subjected to structure prediction from sequence.

## 2 Training on artificial data

The fact that most known RNA structures are derived from a very small set of RNA families makes it hard to distinguish between shortcomings due to the biased training data and more fundamental problems in deep learning for RNA structures. Therefore, we propose to test deep learning methods on completely synthetic data sets generated by classical energy directed structure prediction methods. First, this allows us to test the ability of a neural network (NN) architecture to learn a well-defined biophysical model and discard models that fail to learn certain aspects or are not efficient in doing so. Second, future work can use models that have been pre-trained on synthetic data. This avoids using precious known RNA secondary structures to learn the full complexity of RNA folding from scratch. Instead, a NN that has been trained on the simplified model can subsequently be be presented with known RNA secondary structures to improve on the subtle details of real-world biophysics.

In this contribution we use RNAfold from the ViennaRNA package [10] to fold random sequences allowing us to generate arbitrary large data sets and guarantee complete independence of all sequences in training and evaluation data sets. While structure predictions are far from perfect, they are well known to be statistically similar to the experimentally determined structures [7]. Most results shown below use a training set consisting of random sequences (equal A,U,C,G content) with a homogeneous length of 70 nt, but we also constructed further data sets with four different length distributions, as well as a dataset with the same structure composition as the bpRNA dataset but different sequences. These synthetic sequences enable us to study scenarios where test and evaluation set follow different length and structure distributions.

## 3 Predicting pairedness

To examine what can and cannot be easily predicted by deep learning approaches, we first consider a simplified problem. Rather than predicting base pairs, we restrict ourselves to predict whether a nucleotide is paired or unpaired, in other words, if the nucleotide, in the context of RNA secondary structure belongs to a helix or a loop region. Since this results in a much smaller structure space, one might expect the prediction problem to become easier to learn. This also corresponds to the traditional approach in protein secondary structure prediction, where each amino acid is predicted to be in one of three states (alpha helix, beta sheet, or coil) while ignoring which residues form hydrogen bonds to each other in a beta sheet. Note also that chemical probing of RNA structures [20] typically yields information on pairedness only.

We implemented three different types of predictors: (i) a simple feed-forward network (FFN) that examines sequence windows and predicts the state of the central residue, (ii) a more complex 1D convolutional neural network (CNN), again working on sequence windows, and (iii) a window-less bi-directional long short term memory (BLSTM) network, see Fig. 1. We tested several window sizes for the sliding window approaches (i and ii) and varied the number of layers and neurons in the BLSTM. The FFN architecture is inspired by classical protein secondary structure predictors, such as PHD [16].

**Figure 1:**
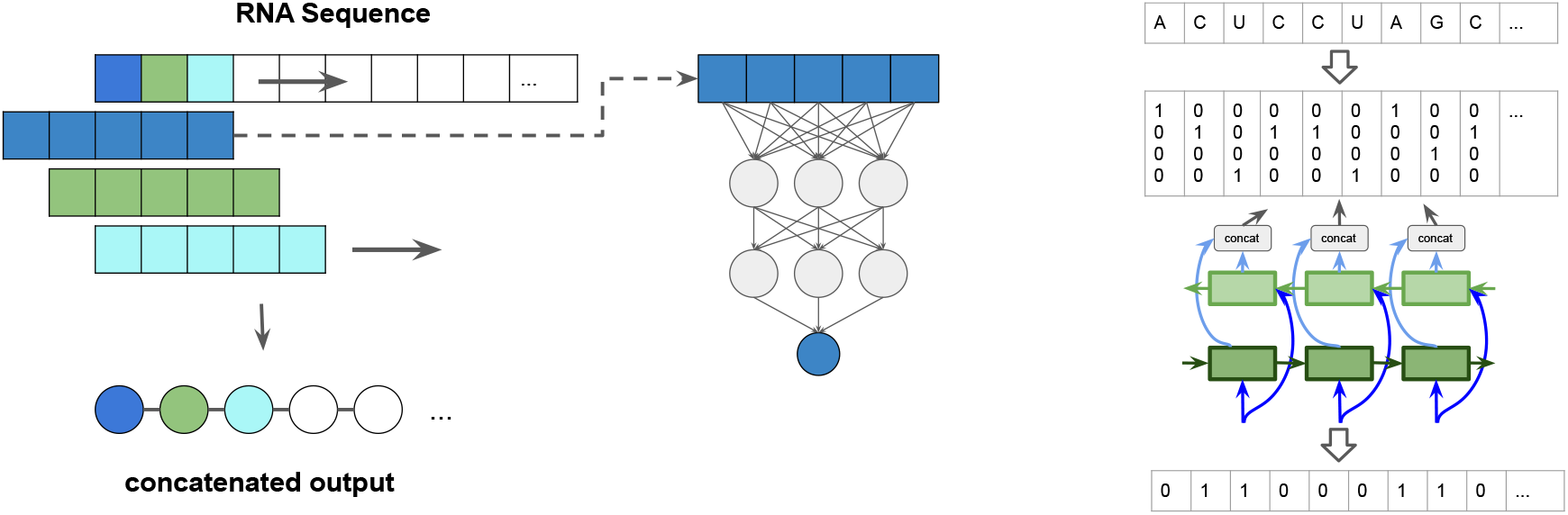
Paired / unpaired prediction approach: (left) sliding-window: A window, consisting of a central symbol and context in the form of a fixed number of leading and tailing symbols is slid along the sequence. The output sequence is a concatenation of the single predictions per window position. (right) schematic representation of the input / output encoding for the bidirectional long short term memory (BLSTM) neural network. The detailed network architectures can be seen in Fig. S2.

The resulting performance when training on sequences of length 70 is shown in Table 1. While the BLSTM performed slightly better than the simpler sliding window approaches, none of the predictors achieved satisfactory performance. This is most obvious when focusing on the Matthews Correlation Coefficient (MCC) [4]. For this task, an accuracy of 0.5 corresponds to pure chance and thus the networks did little more than learn that “A” nucleotides have a higher propensity to be unpaired than “G”s. Our results also indicate that the performance does not improve by increasing the number of neurons, or by using more training data (results not shown). The results were also consistent for different datasets and different training runs.

**Table 1:**
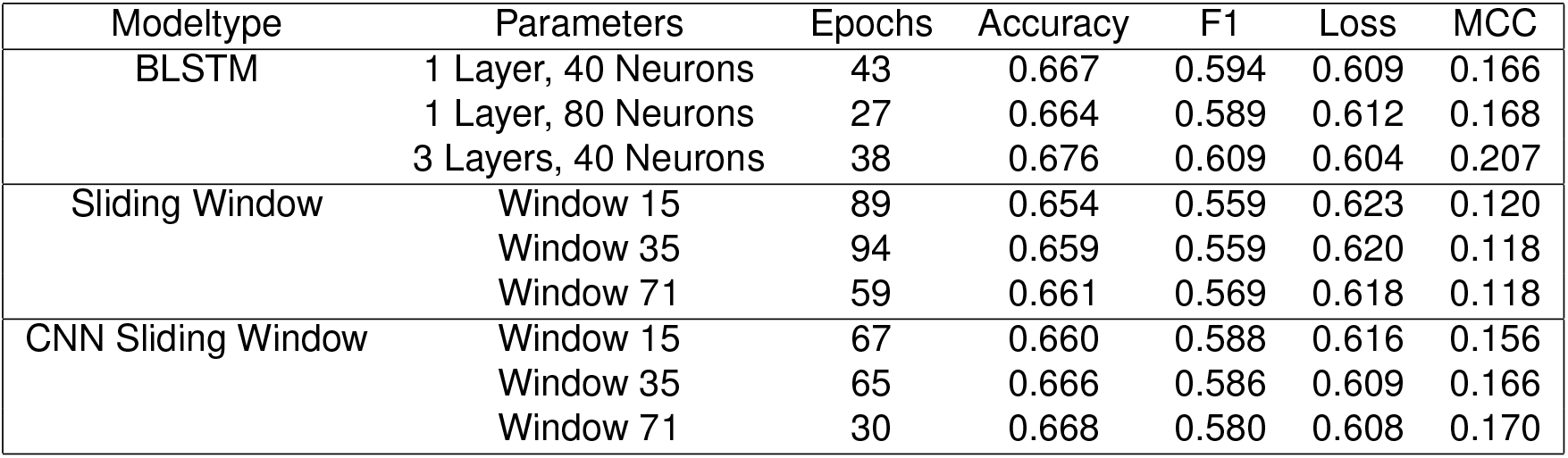
Performance of the paired / unpaired prediction: The performances on the validation set of 20000 sequences of length 70 for all models trained on 80000 sequences of length 70 for 100 epochs. After 100 epochs the best performing model is chosen based on maximum validation MCC. The epoch in which this performance is reached can also be seen in the table. The metrics used are accuracy, F1, loss and MCC. All values are rounded to three decimal places.

| Modeltype | Parameters | Epochs | Accuracy | F1 | Loss | MCC |
| --- | --- | --- | --- | --- | --- | --- |
| BLSTM | 1 Layer, 40 Neurons | 43 | 0.667 | 0.594 | 0.609 | 0.166 |
|  | 1 Layer, 80 Neurons | 27 | 0.664 | 0.589 | 0.612 | 0.168 |
|  | 3 Layers, 40 Neurons | 38 | 0.676 | 0.609 | 0.604 | 0.207 |
| Sliding Window | Window 15 | 89 | 0.654 | 0.559 | 0.623 | 0.120 |
|  | Window 35 | 94 | 0.659 | 0.559 | 0.620 | 0.118 |
|  | Window 71 | 59 | 0.661 | 0.569 | 0.618 | 0.118 |
| CNN Sliding Window | Window 15 | 67 | 0.660 | 0.588 | 0.616 | 0.156 |
|  | Window 35 | 65 | 0.666 | 0.586 | 0.609 | 0.166 |
|  | Window 71 | 30 | 0.668 | 0.580 | 0.608 | 0.170 |

The poor performance suggests that the short cut simply does not work. While it is possible that attention based models such as transformers would performed a little better, the most likely interpretation is that pairedness cannot be predicted independently of the full secondary structure. Moreover, RNA secondary structure is apparently too non-local for sliding window approaches to succeed. This is also in contrast to the fact that RNA secondary structure formation is thought to be largely independent of tertiary structure.

## 4 Predicting base pair matrices

To account for the non-locality of secondary structure, recent deep learning approaches for RNA secondary structure have focused on predicting base pairing matrices. In the typical approach a sequence of length *n* is expanded to a *n* × *n* matrix, where each entry corresponds to a possible base pair. Convolutional networks (or variants thereof) are then used to predict an output matrix containing the predicted pairs, i.e. a 1 in row *i* and column *j* indicates that nucleotides *i* and *j* form a pair. Various postprocessing steps can be appended to derive a valid secondary structure from the pair matrix. Since we were interested in analyzing the performance of the network, we avoided any sophisticated postprocessing and either directly analyses the output matrix (with values between 0 and 1), or obtained a single secondary structure by retaining only the highest entry per row, rounding to obtain values of 0 or 1, and removing pseudoknots.

For our experiments we re-implemented the SPOT-RNA network [18], a deep network employing ResNets (residual networks), fully connected layers and 2D BLSTMs, see Fig. S3. The paper explored several variants, that differ in the size or presence of the different blocks. Most experiments were performed on 3 models, corresponding to Models 0, 1, and 3, in the SPOT-RNA paper. Of these, Model 3 is the only on containing the BLSTM block.

We first tested the simple scenario where all sequences in the training and evaluation sets have the same length of 70 nt. The three models achieved a performance in terms of MCC of 0.554 for model 0, 0.580 for model 1 and 0.640 for model 3. This is quite similar to the values reported for SPOT-RNA after initial training, though models 0 and 1 perform slightly worse in our case. Since model 3 (with BLSTM block) had the best overall performance in this test, and since all three models exhibited very similar behavior in all experiments, we will show only results for Model 3 in the following.

The bpRNA data set shows a very uneven distribution of sequence lengths, with most sequences in the range of 70-120 nt, the length of tRNAs and 5S rRNAs (see Fig S1). We therefore explored scenaria where the length distribution of sequences in test and training set differs, by generating four synthetic data sets with sequences of 25–100 nt, but markedly different length distributions, see Fig. 2. In each case, the training set consisted of 30000 and the validation set 5000 different random sequences.

**Figure 2:**
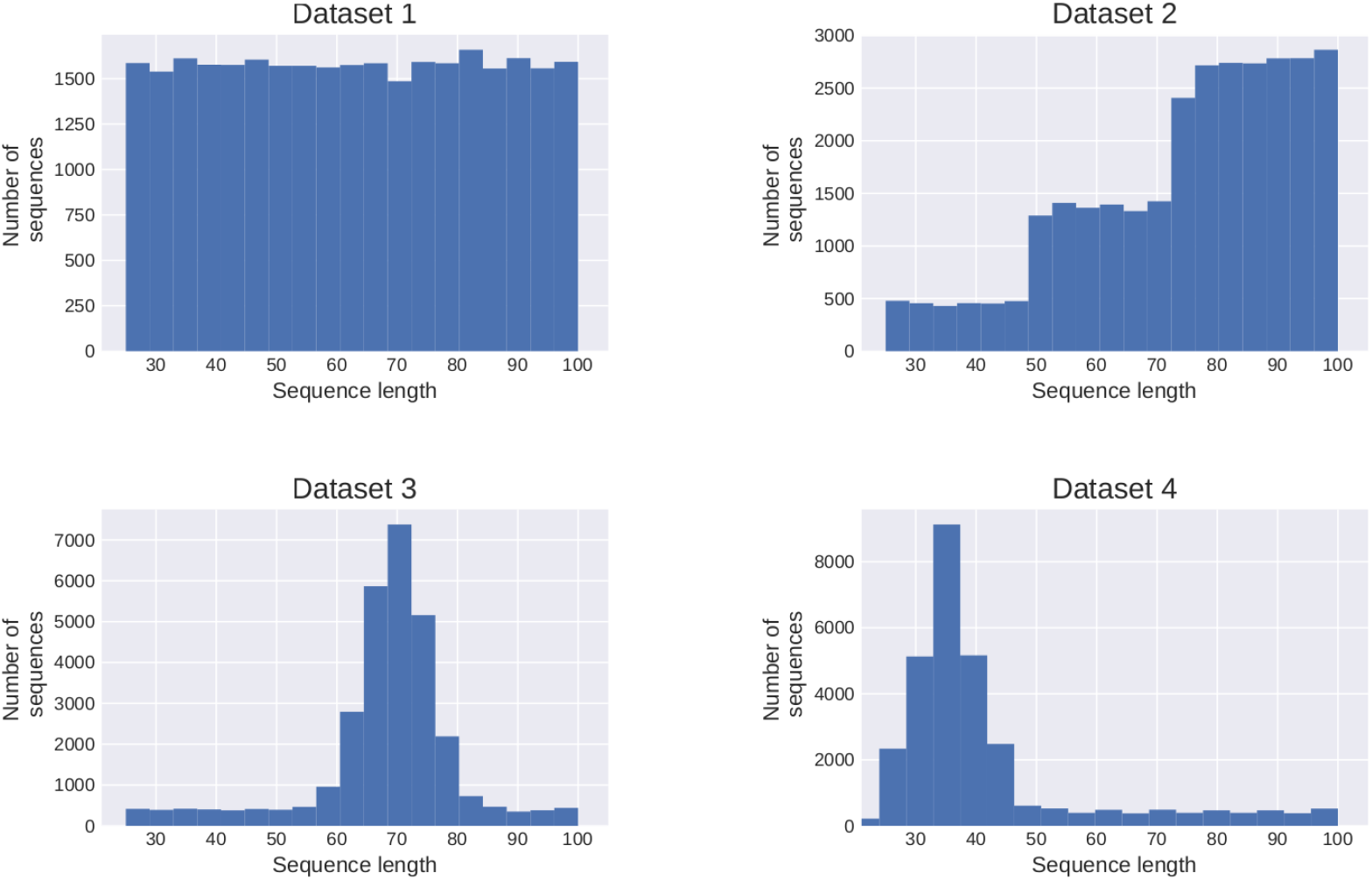
Length distribution of the four synthetic datasets used for prediction of base pair matrices.

We then trained and evaluated our networks on all 16 combinations of training and evaluation sets. Results for Model 3 are shown in Table 2. Even though the datasets were restricted to a rather small range of lengths, from 25 to 100 bases, notable differences are already observable. In general, performance on validation set 4 is best, simply because it contains mostly very short sequences whose structures are easier to predict. Conversely, networks trained on set 4 perform poorly on longer sequences. In addition, we usually observe better performance when training and evaluation set follow the same length distribution, as seen in the diagonal entries of the table. This happens even though all sets are perfectly independent.

**Table 2:** The performances of all combinations of training and validation data sets for the four distributions shown in Figure 2. The diagonal in red shows the performance, when training and validation dataset have the same distribution.

| Training set | Validation set |  |  |  | Performance<br>(training set) |
| --- | --- | --- | --- | --- | --- |
|  | 1 | 2 | 3 | 4 |  |
| 1 | **0.64** | 0.59 | 0.61 | 0.71 | 0.72 |
| 2 | 0.61 | **0.58** | 0.59 | 0.68 | 0.66 |
| 3 | 0.64 | 0.60 | **0.62** | 0.70 | 0.71 |
| 4 | 0.63 | 0.57 | 0.59 | **0.75** | 0.87 |

To further analyze how predictions change with sequence length we generated a series of evaluation sets, varying sequence length from 30 to 250 nt. The number of base pairs is expected to grow linearly with sequence length, since a structure of length *n* must form less than *n*/2 pairs. The ground truth provided by RNAfold perfectly follows the expected behavior. However, for all three network models the number of base pairs, as measured by the number of entries in the output matrix > 0.5 (i.e. before any postprocessing), grows quadratically (see Fig. 3). This happens, because the output matrix has *n*^2^ entries and, asymptotically, the networks predict a constant fraction of all possible base pairs.

**Figure 3:**
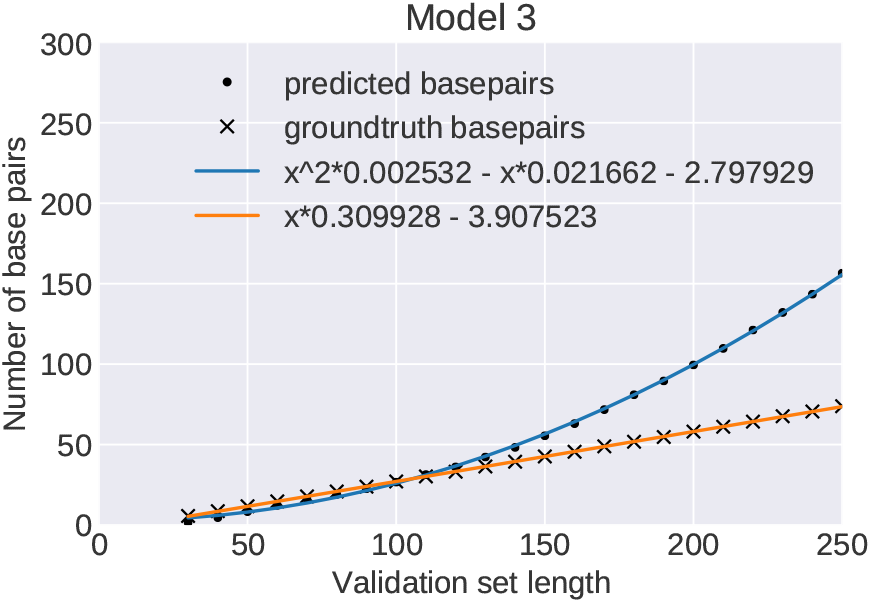
Predicted number of base pairs: Average number of base pairs predicted by model 3 (bullets) and in the ground truth data set (crosses) for 2000 sequence per length bin (30-250). The blue and orange curves are least-square regression fits of the data points. The ML-model predicts a wrong quadratic growth (blue curve) for the number of base pairs in contrast to a correct linear growth (orange line). Results for Models 0 and 1 are indistinguishable

This failure of the network models to reproduce the correct asymptotic behavior exemplifies that it is much easier to learn local properties than global ones. We therefore compared the statistics for several additional structural properties between NN predicted structures and the RNAfold ground truth.

As can be seen in Table 3, the network almost perfectly recapitulates the relative frequency of GC vs AU vs GU pairs and essentially never predicts non-canonical pairs. Frequency and length of hairpin and interior loops are learned quite well. The largest discrepancy is observed for multi-loops, where the network predicts more nucleotides in multi-loops even though it predicts fewer such loops. Consequently, the median length of multi-loops at sequence length 100 is 9 for RNAfold and 16 for model 3. The full distribution of multi-loop lengths is shown in Fig S4. Multi-loops are, of course, harder to learn since they are rarer than the other types and also less local.

**Table 3:** Predicted structural features for RNAfold (VRNA) and Model 3 (NN) trained on sequences of length 70. The test sets consisted of 2000 sequences each of lengths 70 and 100.

| Frequency of bases in context<br>external loop (EL), bulge loop (BL), hairpin loop (HL), internal loop (IL), multi loop (ML), base pairs (bps) |  |  |  |  |  |  |  |
| --- | --- | --- | --- | --- | --- | --- | --- |
| model / length | paired | EL | BL | HL | IL | ML |  |
| VRNA / 70 | 0.508 | 0.176 | 0.033 | 0.156 | 0.114 | 0.014 |  |
| NN / 70 | 0.445 | 0.222 | 0.027 | 0.161 | 0.127 | 0.019 |  |
| VRNA / 100 | 0.541 | 0.123 | 0.031 | 0.143 | 0.126 | 0.035 |  |
| NN / 100 | 0.433 | 0.185 | 0.030 | 0.146 | 0.152 | 0.053 |  |
| Average number of structural element |  |  |  |  |  |  |  |
| model / length | helix | EL | BL | HL | IL | ML |  |
| VRNA / 70 | 4.825 | 0.992 | 1.112 | 1.754 | 1.841 | 0.118 |  |
| NN / 70 | 4.354 | 0.993 | 0.840 | 1.730 | 1.686 | 0.098 |  |
| VRNA / 100 | 7.132 | 0.991 | 1.586 | 2.314 | 2.889 | 0.343 |  |
| NN / 100 | 6.146 | 0.991 | 1.080 | 2.135 | 2.632 | 0.299 |  |
| Relative frequency of base pair types) |  |  |  |  |  |  |  |
| model / length | GC | CG | AU | UA | GU | UG | NC |
| VRNA / 70 | 0.257 | 0.262 | 0.169 | 0.170 | 0.071 | 0.071 | 0.00 |
| NN / 70 | 0.258 | 0.260 | 0.170 | 0.172 | 0.070 | 0.070 | $9.63 \cdot 10^{-5}$ |
| VRNA / 100 | 0.262 | 0.255 | 0.173 | 0.170 | 0.068 | 0.071 | 0.00 |
| NN / 100 | 0.257 | 0.252 | 0.177 | 0.175 | 0.068 | 0.070 | $2.30 \cdot 10^{-5}$ |

A noteworthy effect of learning from data generated by the Turner model is that the training data does not contain pseudoknots or bases forming more than one pairs. However, even though no pseudoknots were available in the training set, out of 2000 structures with length 100, 975 structures contained one or more non-nested base-pairs (2407 base-pairs total), and 1512 structures contained one or more nucleotide with multiple pairs (6250 multi-pairs total). The ability to predict pseudoknots is an attractive feature of neural networks. However, since networks predict pseudoknots, even if the ground truth is pseudoknots free, this casts doubt on the networks ability to learn which potential pseudoknots will form in reality.

## 5 The Effect of biased Training Data

As mentioned earlier the commonly used bpRNA is heavily biased towards a small number of RNA families and thus exhibits little structural diversity. To explore the effect of this bias, we generated an artificial training set with the exact same bias. We used each structure contained in bpRNA and generated a sequence folding into this structure using the RNAinverse program of the ViennaRNA package. If RNAinverse was not able to generate a sequence folding exactly into the target structure, the best sequence out of 6 RNAinverse runs was used. As before, the synthetic data use the RNAfold predicted structure as ground truth. While it is maximally diverse with respect to sequences, this bRNAinv data set carries the same structure bias as the original bpRNA data set. For efficiency reasons, we only used RNAs with a length between 25 and 120 nt. Furthermore, we removed pseudoknots from the bpRNA structure and eliminated structures containing more than 6 base pairs in pseudoknots. This resulted in a data set of 65766 sequence/structure pairs that was further split into training and validation sets of 52613 and 13153 sequences, respectively.

We now trained our Model 3 on this data set. We started from a network that had been pre-trained for just 3 epochs on random sequences of length 119 and 120, since we had noted that such pre-training led to faster training, and then performed 40 epochs of training on the inverse folded bpRNA training set. The model eventually achieved an MCC of over 0.9 on the training set and over 0.85 on the validation set.

To test whether the model was able to generalize to RNAs with structures other than those contained in the bpRNA data set, we generated a test set containing 10000 sequences by inverse folding as explained above. This inverse folded dataset has the same structure bias as the bpRNA dataset and the training set. A randomized version of the test set was created by di-nucleotide shuffling each sequence using ushuffle[9] and computing the corresponding structure using RNAfold. This shuffled dataset has the same sequence composition, but a much higher structural diversity. On the inverse folded tests set (again replicating the structure bias of the training set) the network achieved excellent prediction accuracy with an MCC of 0.86, on the shuffled sequences, however, MCC dropped to only 0.52. This effect was not visible on the pre-trained network where both inverse folded RNAs and shuffled RNAs achieved an MCC of 0.46. During training, performance on the inverse dataset improved rapidly within a few epochs, while performance of the suffled dataset improved only marginally.

## 6 Conclusion

The performance of deep networks is strongly dependent on quantity and quality of the available training data. Biological data, however, are strongly biased towards a small number of well studied model systems. This makes it hard to study the capabilities and shortcomings of networks independently of the quality of available data. This problem can be avoided, if there is a way to generate synthetic training data that are statistically sufficiently similar to real data. By performing machine learning experiments on the synthetic data one can identify strengths and shortcomings of different network architectures, determine the amount of training data required, and study the effect of biases in the training data. For RNA secondary structure prediction, algorithms that compute the minimum free energy structure via dynamic programming on a biophysical energy model can provide such a data source.

While recent RNA secondary structure data sets provide a large number of training sequences, this comes at the expense of making the data set extremely unbalanced, with more than 95% of sequences deriving from ribosomal RNAs or tRNAs. Neural networks learn to exploit this bias, leading to predictors that perform very well on RNAs whose structures are well represented in the training set. The networks generalize well to RNAs with no *sequence* similarity, as long as the structure has been seen in the training set. However, the performance on new structures remains poor. Even on otherwise unbiased synthetic training sets, performance suffers when training set and evaluation set follow a different length distribution. These results emphasize the importance of using training and test sets that are structurally distinct, i.e. no RNA family should be present in both training and test set. An ideal data sets should fairly sample the space of secondary structures, but this would require large scale structure determination beyond selected biologically interesting RNAs.

Our experiments with training on synthetic data also reveal which properties of RNA structure are easy or hard to learn for current network architectures. Features such as base pairs, interior-, and hairpin-loops, are local with respect to the base pairing matrix and easy to learn. Indeed, the prevalence of different types of base pairs, as well as length and size of hairpin and interior loops, almost perfectly matches the ground truth. Multi-loops and pseudoknots are less local and exhibit significant deviations from the ground truth. Finally, the networks struggle to correctly reproduce global properties and scaling behavior, as exemplified by the fact that for all networks the number of predicted base pairs scales quadratically with sequence length. While this behavior can easily be addressed during postprocessing, it is not clear whether that would correct or merely hide the underlying problem.

## Author Contributions

ILH and CF conceived and supervised the study. JW implemented the machine learning models, trained models on synthetic data with different length distributions and performed the initial statistical analysis. SB and ILH trained and analyzed the models on inverse bpRNA data. SB and MTW helped with statistical data analysis. MTW and RL were involved in planing and supervision of the machine learning experiments. ILH and CF wrote the first draft of the manuscript. All authors contributed actively to rewriting the draft into the final manuscript. The final manuscript has been read and approved by all authors.

## Acknowledgments

This is a short text to acknowledge the contributions of specific colleagues, institutions, or agencies that aided the efforts of the authors.

## Supplemental Data

Additional figures with details on the neural network architectures can be found in the supplement.

## Data Availability Statement

Python notebooks implementing our models and data files are available at https://github.com/ViennaRNA/RNAdeep

