## Supplementary data for "Caveats to deep learning approaches to RNA secondary structure prediction"

### Supplementary Material

#### 1 SUPPLEMENTARY DATA

The ML-modles and the trainingsdata can be found on github `https://github.com/ViennaRNA/RNAdeep`

#### 2 SUPPLEMENTARY TABLES AND FIGURES

##### 2.1 Figures

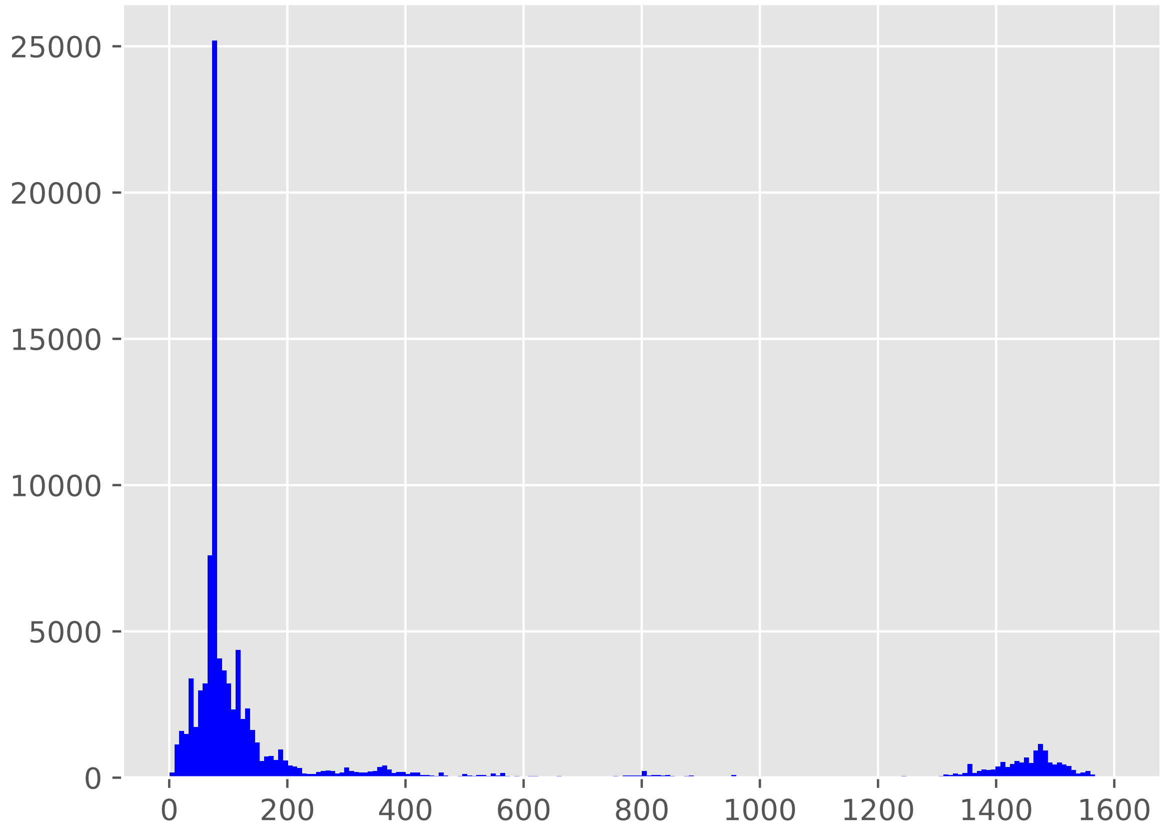


**Figure S1.** Length distribution of the sequences in the bpRNA-1m dataset Version 1 `http://bprna.cgrb.oregonstate.edu/`. The highest peak correspond to tRNAs of a length of about 75 nucleotides (nts). For the plot the dataset was truncated at length 1600 nts (removing 736 sequences longer than 1600 nts).

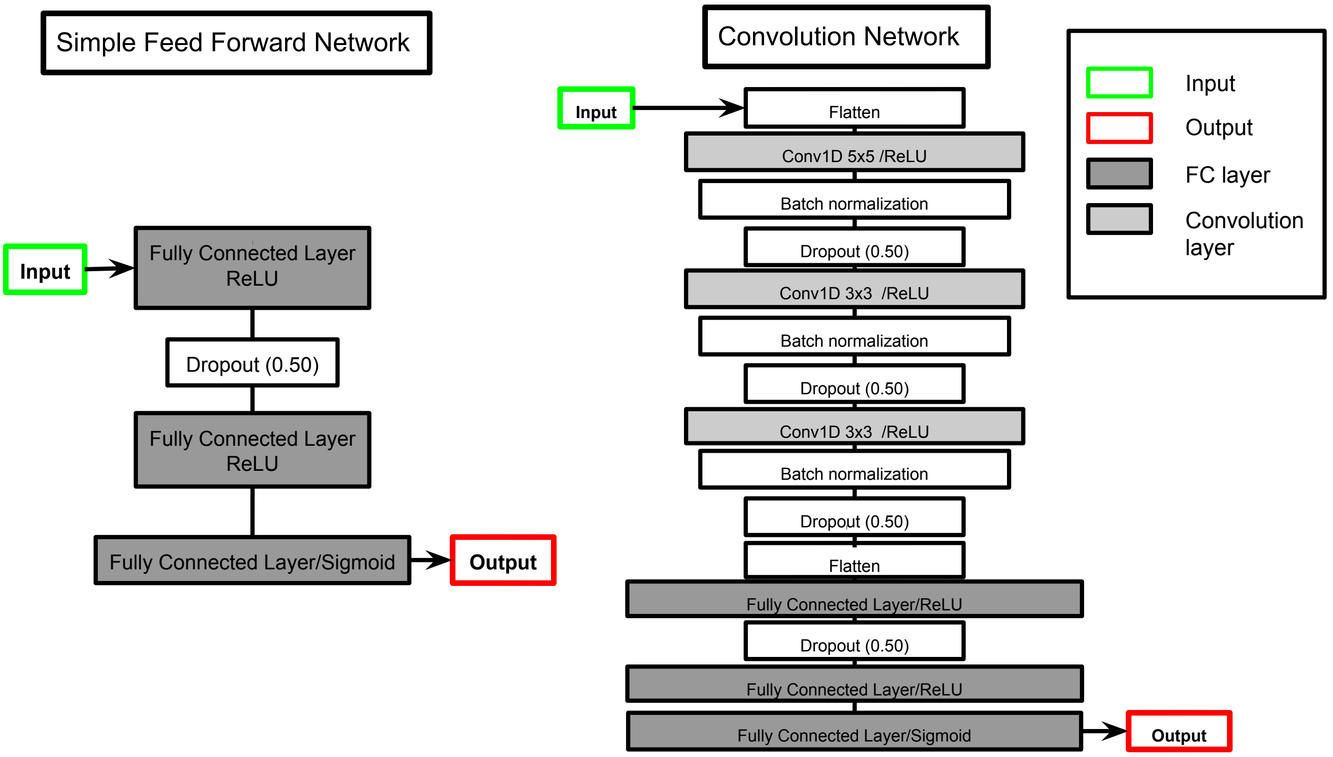


**Figure S2.** Neural network architectures for the sliding window approach. (left) feed forward neural (FFN) network (right) 1D convolutional neural network (CNN)

doi:10.1038/s41467-019-13395-9

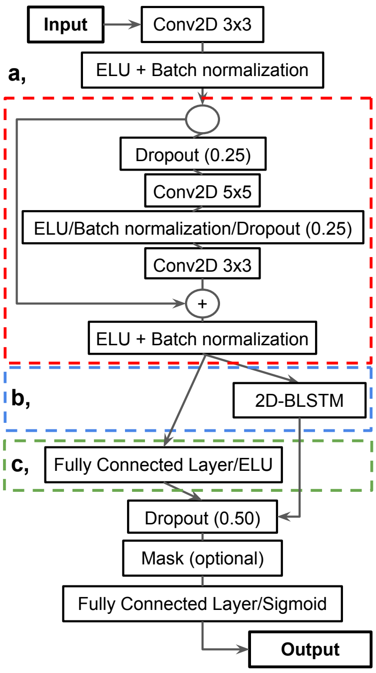


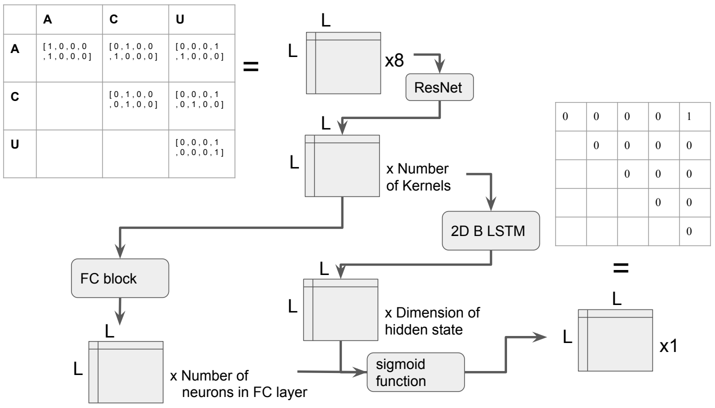


| Model | Filters | ResNet blocks | LSTM blocks | LSTM neurons | FC blocks | FC neurons |
|---|---|---|---|---|---|---|
| 0 | 64 | 20 | 0 | 0 | 2 | 512 |
| 1 | 48 | 16 | 0 | 0 | 2 | 512 |
| 3 | 64 | 30 | 1 | 200 | 0 | 0 |

**Figure S3.** Detailed view of the reimiplemented network architectures of the SPOT-RNA network (Singh et al., 2019) refered to as model 0-3. The repeat numbers for the ResNet (in red), the LSTM (in blue) and the fully connected layers (in green) are listed in table on the bottom right. How the data is fed into and retreved from the models is shown in the schematic figure on the top right.

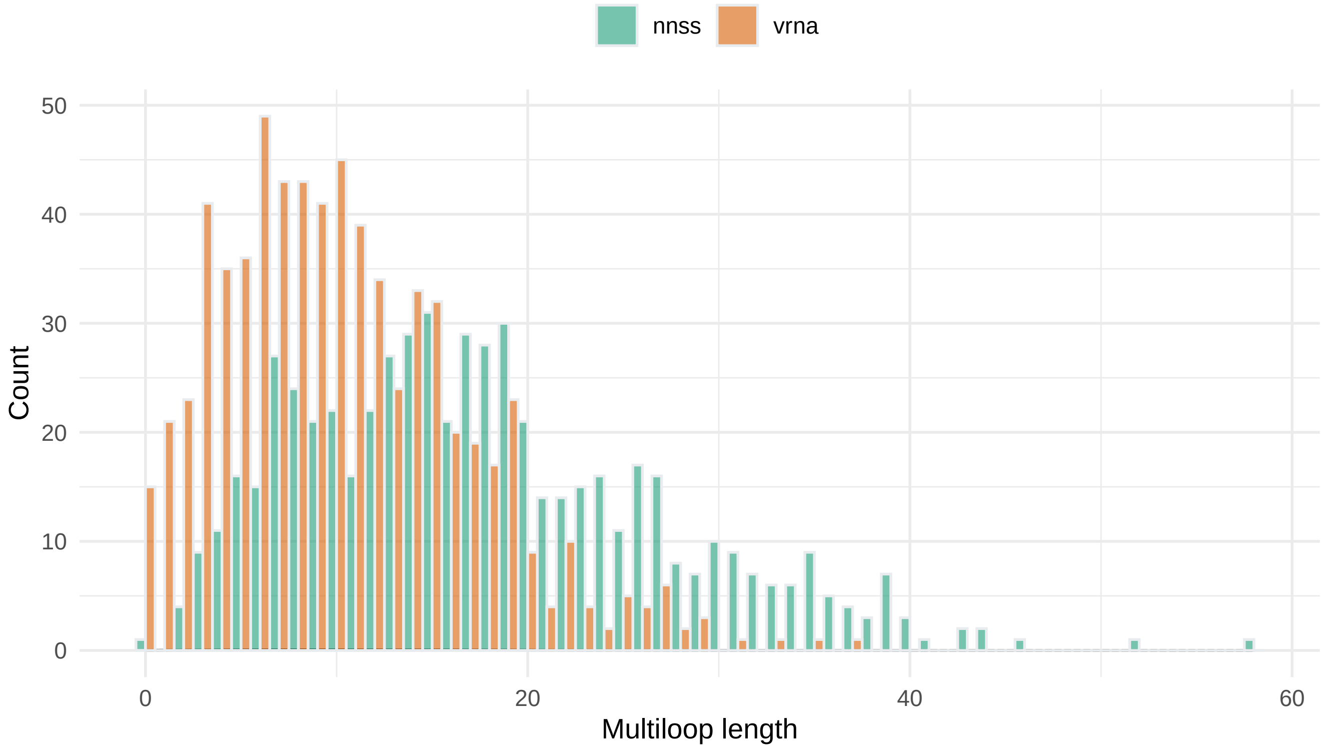


**Figure S4.** Length distribution of multi-loops of the validation set of length 100 nt predicted by the neural network (nnss) and `ViennaRNA` (vrna), respectively.
